# Echidna: integrated simulations of single-cell immune receptor repertoires and transcriptomes

**DOI:** 10.1101/2021.07.17.452792

**Authors:** Jiami Han, Raphael Kuhn, Chrysa Papadopoulou, Andreas Agrafiotis, Victor Kreiner, Danielle Shlesinger, Raphael Dizerens, Kai-Lin Hong, Cédric Weber, Victor Greiff, Annette Oxenius, Sai T. Reddy, Alexander Yermanos

**Affiliations:** Department of Biosystems Science and Engineering, ETH Zurich, Basel, Switzerland; Institute of Microbiology, ETH Zurich, Zurich, Switzerland; Department of Pathology and Immunology, University of Geneva, Geneva, Switzerland; Department of Immunology, University of Oslo, Norway

## Abstract

Single-cell sequencing now enables the recovery of full-length immune repertoires [B cell receptor (BCR) and T cell receptor (TCR) repertoires], in addition to gene expression information. The feature-rich datasets produced from such experiments require extensive and diverse computational analyses, each of which can significantly influence the downstream immunological interpretations, such as clonal selection and expansion. Simulations produce validated standard datasets, where the underlying generative model can be precisely defined and furthermore perturbed to investigate specific questions of interest. Currently, there is no tool that can be used to simulate a comprehensive ground truth single-cell dataset that incorporates both immune receptor repertoires and gene expression. Therefore, we developed Echidna, an R package that simulates immune receptors and transcriptomes at single-cell resolution. Our simulation tool generates annotated single-cell sequencing data with user-tunable parameters controlling a wide range of features such as clonal expansion, germline gene usage, somatic hypermutation, and transcriptional phenotypes. Echidna can additionally simulate time-resolved B cell evolution, producing mutational networks with complex selection histories incorporating class-switching and B cell subtype information. Finally, we demonstrate the benchmarking potential of Echidna by simulating clonal lineages and comparing the known simulated networks with those inferred from only the BCR sequences as input. Together, Echidna provides a framework that can incorporate experimental data to simulate single-cell immune repertoires to aid software development and bioinformatic benchmarking of clonotyping, phylogenetics, transcriptomics and machine learning strategies.

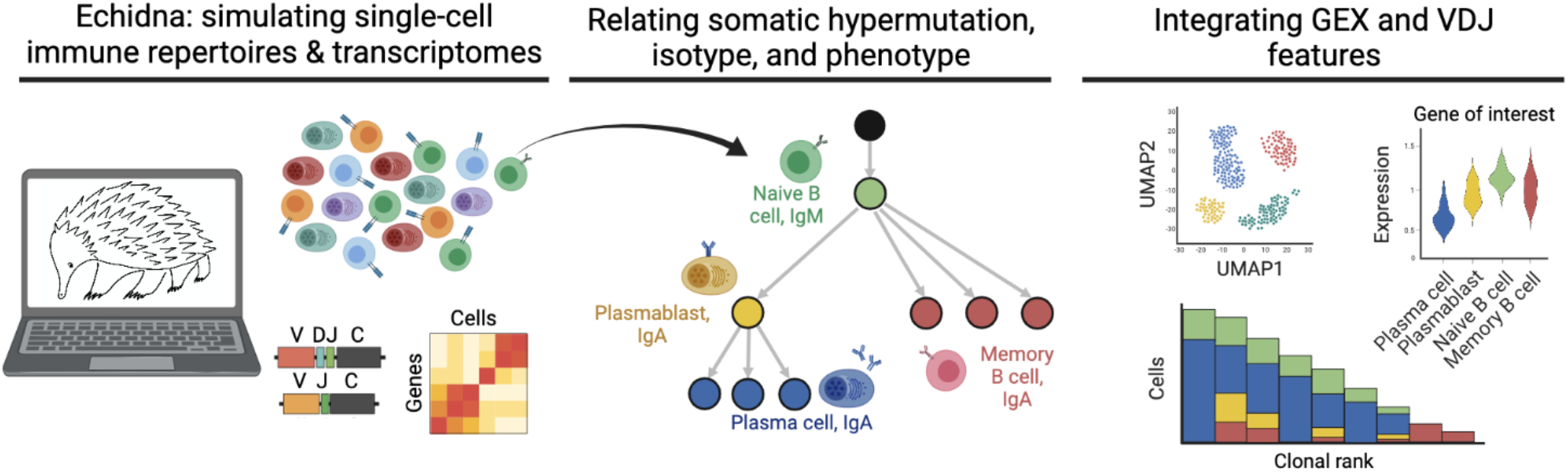

## Introduction

The adaptive immune system plays a crucial role in the protection against a broad range of pathogens. The molecular recognition of foreign pathogens by B and T cells is orchestrated by their characteristic BCRs and TCRs, respectively. Sequence diversification is first introduced into BCRs and TCRs by the recombination of genomically encoded variable (V), diversity (D), and joining (J) gene segments (Tonegawa, 1983). This diversity is further increased by the combinatorial pairing of variable heavy (V_H_) and variable light (V_L_) chains for B cells and variable beta (V_b_) and variable alpha (V_a_) chains for T cells. BCRs and their secreted form, antibodies, can undergo additional diversification through somatic hypermutation (SHM) (Diaz and Casali, 2002). Together, these processes result in a vast diversity (10^18^ and 10^13^ for B and T cells, respectively in humans), which provides the baseline interaction space that accommodates a seemingly infinite number of foreign antigens (Greiff *et al.*, 2017; Murphy and Weaver, 2016).

Advances in deep sequencing have provided the opportunity to quantify and map sequence diversities of immune repertoires. Indeed, repertoire sequencing has been instrumental to investigate fundamental immunological principles in the context of disease, infection, and immunization (Neumeier, Pedrioli, *et al.*, 2021; Horns *et al.*, 2020; Yermanos, Agrafiotis, *et al.*, 2021a). However, prior to the emergence of single-cell sequencing, repertoire sequencing was performed on bulk (pooled) cell samples, which has a number of limitations: inability to recover full-length, paired sequences (V_H_ + V_L_ for B cells and V_b_ + V_a_ for T cells), the reliance upon read counts as a proxy for clonal expansion, and lack of individual cell phenotyping based on surface marker expression (Rosati *et al.*, 2017; Yaari and Kleinstein, 2015; Miho *et al.*, 2018; Georgiou *et al.*, 2014; Brown *et al.*, 2019). The advent of single-cell sequencing (scSeq) has revolutionized the resolution at which clonal selection of immune repertoires can be quantified (Friedensohn *et al.*, 2017). For example, scSeq workflows have been established that enable the simultaneous recovery of both full-length, paired BCR or TCR sequence (VDJ), and full transcriptome [gene expression (GEX)] information, thereby providing a crucial link between clonal selection and expansion with a high-dimensional transcriptional cell phenotype (Horns *et al.*, 2020; Khatun *et al.*, 2021; Yermanos, Neumeier, *et al.*, 2021; Mathew *et al.*, 2021; Lindeman *et al.*, 2018; Stubbington *et al.*, 2016). Interpreting such datasets, however, remains challenging as the accompanying computational pipelines and software are still in their infancy (Yermanos, Agrafiotis, *et al.*, 2021b; Borcherding *et al.*, 2020; Sturm *et al.*, 2020). Although multiple tools have been developed to simulate immune receptors and single-cell transcriptomes (Marcou *et al.*, 2018; Weber *et al.*, 2020; Yermanos *et al.*, 2017; Davidsen *et al.*, 2019; Safonova *et al.*, 2015), these represent separate platforms and thus there remains a lack of software capable of simulating scSeq data of immune repertoires and transcriptomes. We therefore developed Echidna, an R package that simulates immune repertoires and their corresponding transcriptomes. Importantly, datasets are constructed in a style compatible with commonly used methods of scSeq (e.g., 10X Genomics). We furthermore demonstrated the applicability of Echidna by simulating immune repertoires with varying features such as cell phenotype, clonal expansion, somatic hypermutation, antibody isotype, and germline gene usage. Finally, we leveraged Echidna simulated and time-resolved B cell mutational networks and evaluated the accuracy when using clonal lineage reconstruction algorithms.

## Results

### Echidna simulates time-resolved single-cell immune receptor repertoires

To address the lack of a software capable of simulating integrated immune receptors and transcriptomes, we developed the R package Echidna. Echidna generates full-length and paired adaptive immune receptors with the corresponding gene expression information at single-cell resolution for both human and murine immune repertoires (Figure 1). Importantly, the format of the resulting simulation mirrors the output files of the widely used 10x Genomics cellranger tool (Figure S1), thereby rendering simulated data compatible with existing bioinformatics tools such as Platypus and Seurat (Yermanos, Agrafiotis, *et al.*, 2021b; Satija *et al.*, 2015).

**Figure 1.**
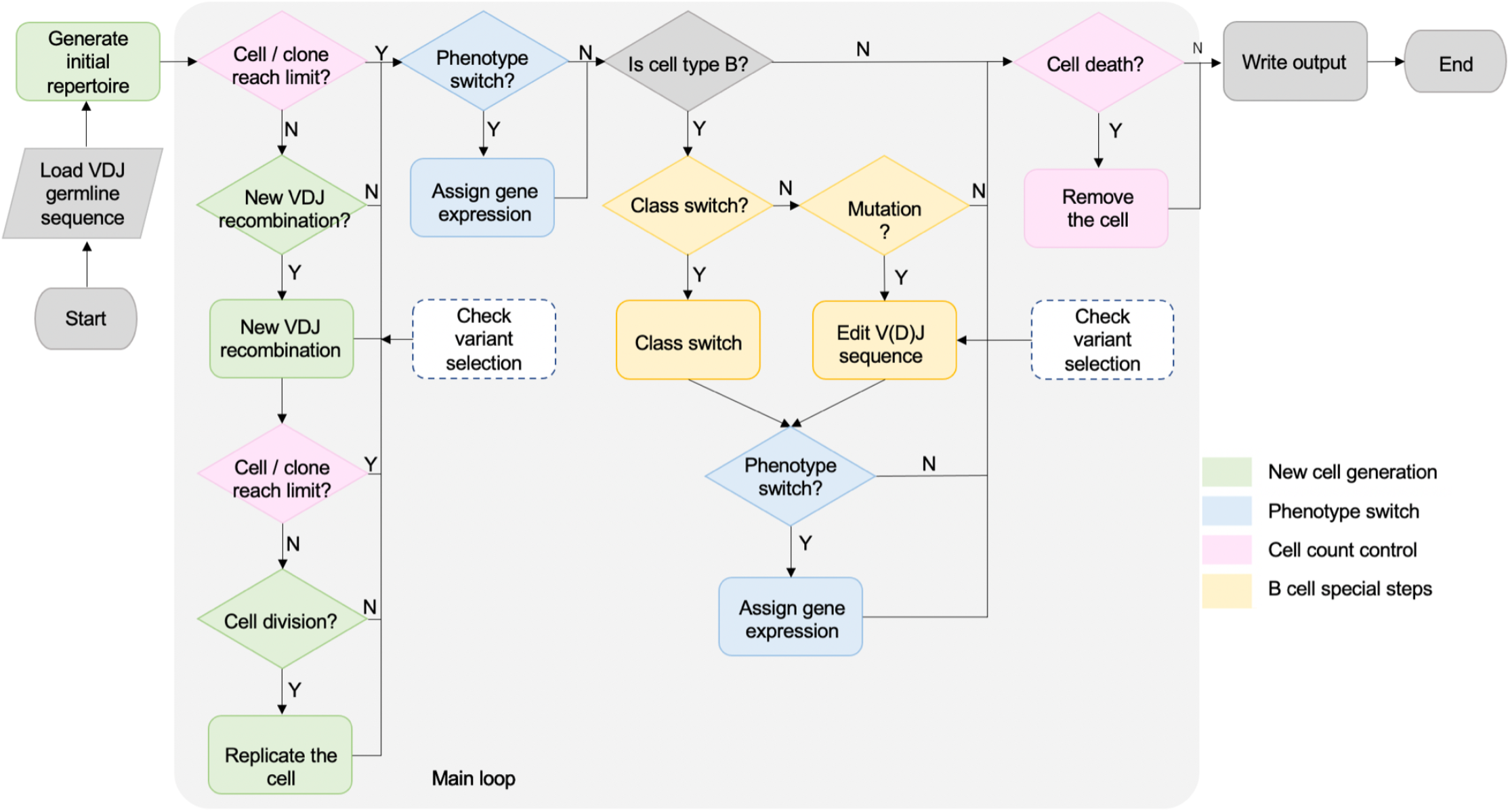
Computational workflow of single-cell immune repertoire and transcriptome simulations using Echidna.

The simulation first generates an initial repertoire of cells, each produced by a separate recombination event for both VDJ (V_H_ for B cells, V_b_ for T cells) and VJ (V_L_ for B cells, V_a_ for T cells) genes. This can either be done as previously described (Yermanos *et al.*, 2017) by appending reference germline segments from the Immunogenetics Database (IMGT) (Lefranc *et al.*, 2003) with diversity generated by insertions and deletions in the complementarity determining region 3 (CDR3) or by generating sequences using a variational autoencoder (Friedensohn *et al.*, 2020) using publicly available BCR and TCR sequences for humans and mice (Kovaltsuk *et al.*, 2018). The user can further specify whether these simulated initial sequences should be restricted to productive BCR and TCR sequences based on alignment and annotations (Bolotin *et al.*, 2015). By specifying the starting repertoire size, which is defined by the number of B or T cell clones at the first simulated time-step, the user can control the initial clonal diversity via simulated V(D)J recombination. Under default parameters, germline genes are uniformly sampled from the IMGT reference database and appended together with a probability of inserting or deleting nucleotides at the junctions (both VD and DJ for V_H_ and V_b_ chains, and VJ for V_L_ and V_a_ chains) following a user-supplied distribution. While under default conditions, each germline gene has a uniform probability of being selected for V(D)J recombination, this distribution can be altered to preferentially utilize certain germline segments as is the case under physiological in vivo settings .

To demonstrate the ability to simulate BCR repertoires, 5,000 V(D)J recombination events were generated in silico where V_H_ and V_L_ genes were selected with a uniform probability distribution or mirrored V-gene distributions observed in experimental studies (Greiff *et al.*, 2017; Kräutler *et al.*, 2020; Yermanos *et al.*, 2020) (Figure 2A). Such a preferential germline gene usage can also be utilized for VJ pairings of either the heavy or light chain, providing the user further control over the initial diversity generated by V(D)J recombination (Figure 2B). In addition to the simplified random assortment of V, D, and J germline genes, one can simulate the starting pool of immune receptors using variational autoencoders (VAEs), which has been recently used to identify convergent, antigen-specific antibodies and for the in silico generation of novel sequences (Friedensohn *et al.*, 2020). While Echidna’s internal VAE will produce productive sequences arising from naive human and murine B and T cell repertoires from either publicly available reference genes (Lefranc *et al.*, 2003) or sequencing data (Kovaltsuk *et al.*, 2018), we have additionally included an internal function that allows the user to supply VDJ or VJ sequences as input to generate novel sequences for an initial repertoire, in case other training datasets are of interest. Both appending reference germlines and VAEs to generate starting VDJ-recombined clones results in a high proportion of unproductive sequences (Figure S2A), which may not be ideal for downstream applications. We, therefore, have added the option to maintain only productive adaptive immune receptor sequences in the initial repertoire for either recombination option (Figures S2B, S2C).

**Figure 2.**
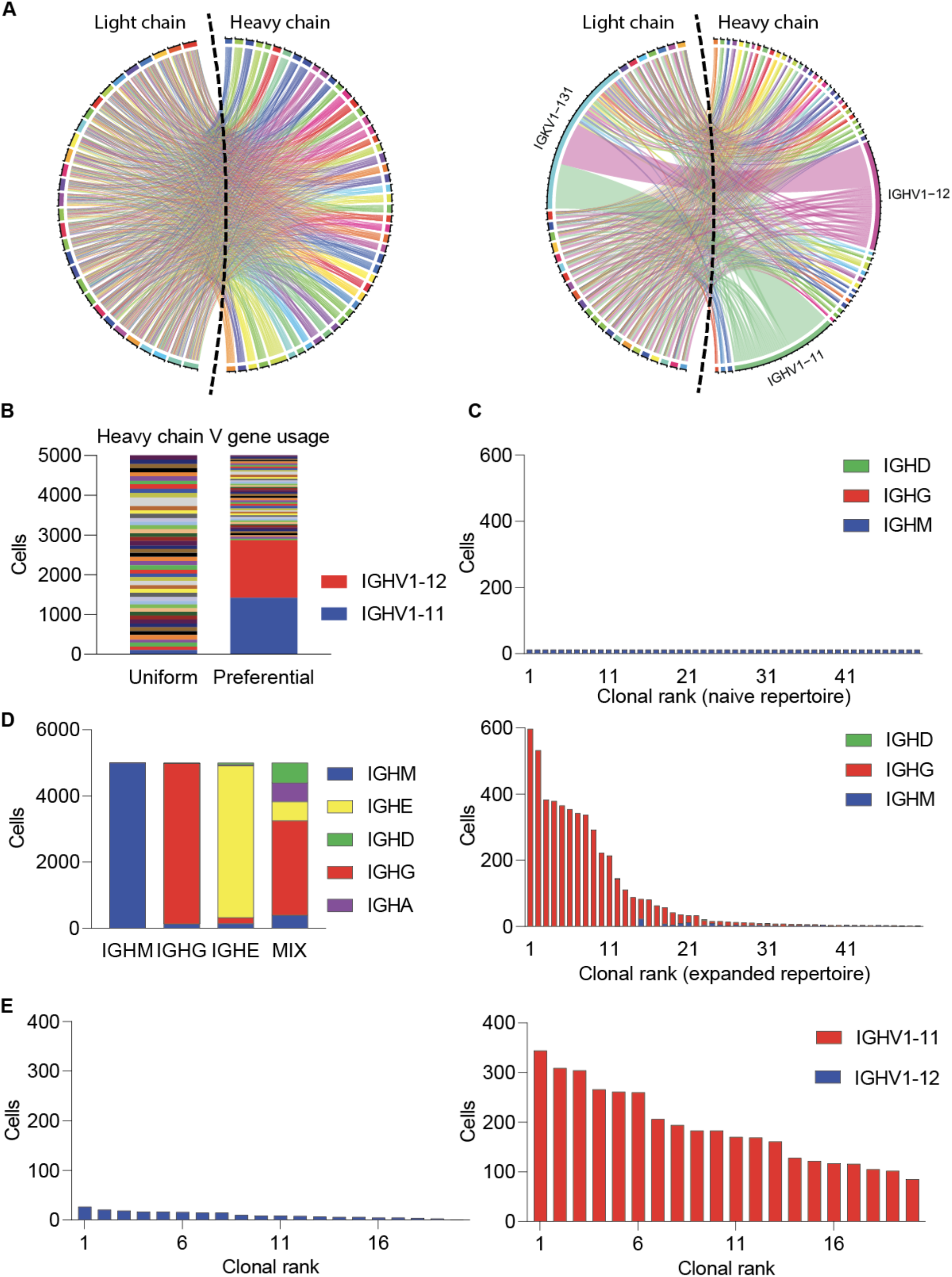
Echidna simulates expanded and naive immune repertoires at single-cell resolution. A. Repertoires with either uniform or preferential germline gene usage of IGHV1-11. Each color represents a unique germline gene. Line width represents the number of cells using a particular variable heavy or variable light chain combination. B. Heavy chain variable gene usage. Color indicates the unique segment. The two preferential germline genes used are specifically highlighted. B. Clonal frequency for either naive-like (top) or expanded (bottom) repertoires. Clone is defined as all cells arising from a single simulated VDJ recombination event. Bar color corresponds to the fraction of cells within each clone of a given isotype. C. Expanded and naive B cell clonal expansion plot of top 50 clones colored by isotype. D. Preferential selection and expansion based on germline gene usage within a single repertoire. Clone is defined as all cells arising from a single simulated VDJ recombination event.

### Repertoire features dictate simulated clonal expansion

Following initialization of the first cell for a given clone (defined by all cells arising from a unique V(D)J recombination event), the time-resolved phase of the simulation begins, where new cells can either be introduced into the repertoire, existing cells can undergo clonal expansion, or cells can be removed from the simulation via cell death. Although the final simulated repertoire does not include the BCRs and TCRs of the cells that have undergone cell death, the history of these cells can still be analyzed and reconstructed by the internal storage of parameters such as adaptive immune receptor sequence. To demonstrate the ability of Echidna to simulate different modes of clonal expansion for B and T cells, we simulated either naive or expanded repertoires by altering the simulation parameters by controlling the probability to either undergo cell division or to recruit new cells into the immune repertoire (Figure 2C). In the case of B cells, cells can additionally undergo class-switch recombination from the starting IgM+ isotype with a user-supplied transition matrix. While the default transition matrix follows a probability distribution mirroring the biological distribution of class-switching (IgM > IgG > IgA > IgE), it is possible to alter this matrix to mirror cases where preferential class-switching occurs, such as viral infections or immunizations (Figure 2D) (Neumeier, Pedrioli, *et al.*, 2021; Neumeier, Yermanos, *et al.*, 2021). While the default settings of Echidna have a uniform probability of cells within a given clone to undergo clonal expansion (and SHM in the case of B cells), the user can further tune this parameter to link particular repertoire features, such as germline gene usage, to clonal expansion, as visualized by certain clones which undergo preferential expansion (Figure 2F). Taken together, these examples demonstrate the ability of Echidna to incorporate repertoire features, such as germline gene usage and isotype, into the model underlying clonal selection.

### Time-resolved evolution of simulated B cell receptors

In the case of B cells, Echidna can simulate SHM and clonal evolution in a time-resolved manner. Specifically, at each step, each cell has a probability to undergo BCR diversification using three distinct mutational models. While the most simple method involves a uniform distribution for each nucleotide to mutate to another base, a second option favors transitions relative to transversions. Finally, a third option is to introduce mutations following an experimentally determined 5-mer mutation model, in which neighboring nucleotides influence transition probabilities (Cui *et al.*, 2016; Yaari *et al.*, 2013). At each simulated time step, each cell has a probability to proliferate and incur SHMs, thereby giving rise to either clonally expanded antibody sequences or variants within an individual clone. While it is possible to modulate the method and number of SHM-derived clonal variants, under default parameters Echidna introduces more mutations to class-switched B cells (Figure 3A). In addition to recovering the full-length antibody sequence associated with each simulated cell barcode, Echidna furthermore provides the mutational network depicting the evolutionary history for each B cell, thereby following the ground-truth of B cell evolution (Figure 3B). The mutational networks contain information pertaining to clonal expansion of individual antibody variants (node size), the fraction of cells with a given isotype (node color), and the clonal relationships (edges). Although both mutations and cell death result in intermediate variants being removed from the final list of output sequences, an additional mutational network including intermediate nodes with their corresponding sequences can be provided (Figure 3C). The information from the mutational network and the output sequences can be integrated to investigate how sequence-level mutations dictate clonal relationships (Figure 3D). To demonstrate Echidna’s ability to benchmark clonal lineage reconstruction algorithms and tools, we simulated 20 B-cell networks and supplied the output sequences and respective germlines into a previously utilized pipeline that creates B cell mutational networks based on converting pairwise edit distances to adjacency matrices (Neumeier, Pedrioli, *et al.*, 2021). Comparing the topologies of the simulated and inferred networks, we could demonstrate that 19 of the 20 simulated networks were correctly predicted, with the major divergence due to the presence of germline-like B cells which closely resemble the unmutated reference germline (Figures 3E, S4).

**Figure 3.**
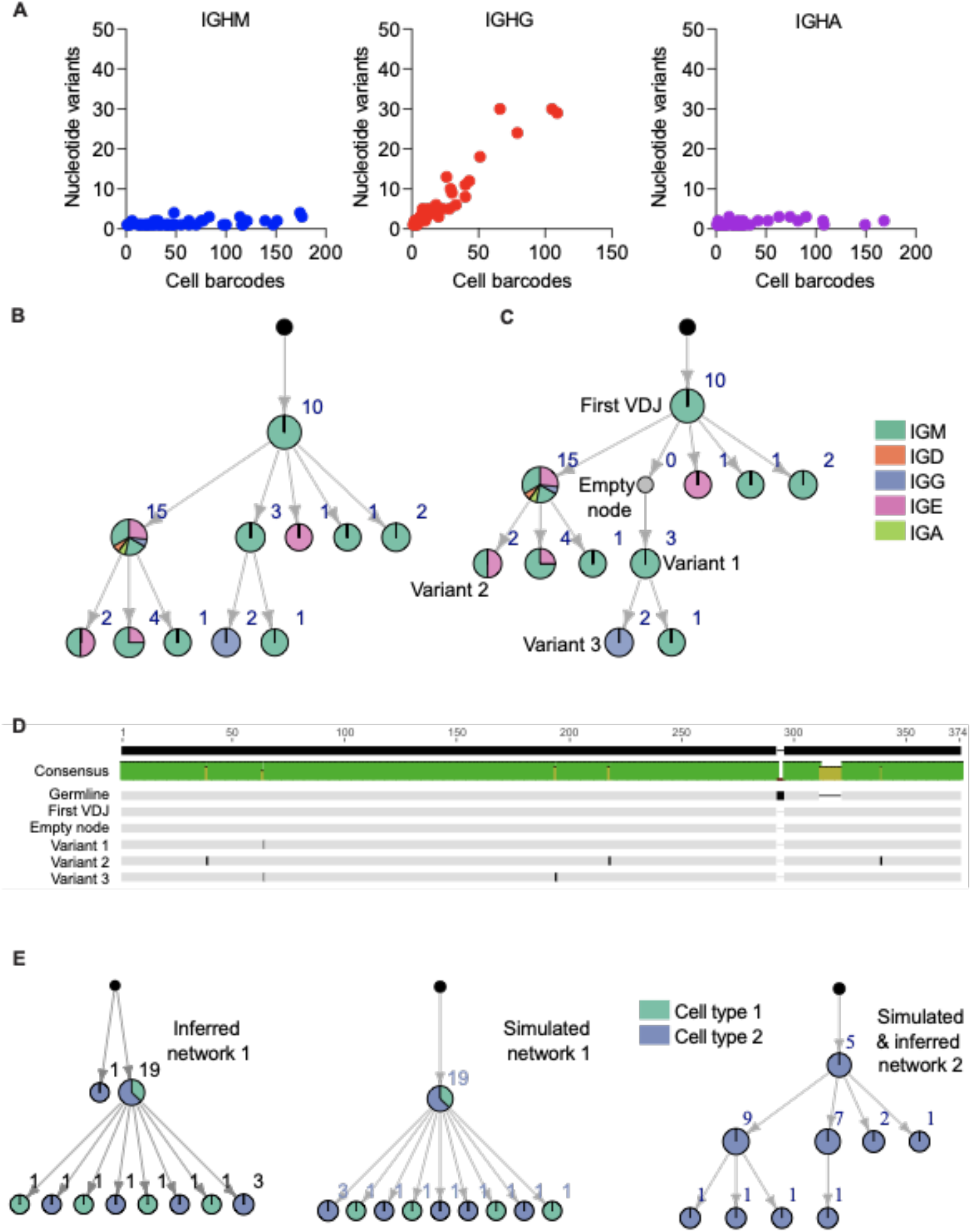
Simulation of time-resolved somatic hypermutation and evolution. A. Relationship between the number of cells and the number of unique BCR sequences for each isotype. Each point represents a clone, defined as all cells belonging to a single simulated VDJ recombination event. Variants are defined by a unique full-length VH-VL nucleotide sequence. B. Simulated clonal lineages depicting the relationship between somatic hypermutation, clonal expansion, and isotype. Each node depicts a unique variant, defined by a unique full-length VH-VL nucleotide sequence. The color of each node corresponds to the proportion of cells within that variant of a certain isotype. Edges depict clonal relationships. Black nodes correspond to unmutated reference germline. C. Clonal lineage as in (B) but including the historical node that is no longer present by the end of the simulation due to mutation or cell death. D. VH sequence alignment from the clonal lineage in C. “Germline” refers to the appended V, D, and J reference genes without additional alterations to the CDR3. “First” refers to the first simulated, productive sequence following the insertion and deletion portions of VDJ recombination. “Empty node” refers to an ancestral sequence that is no longer present in the final simulation. Variants 1, 2, and 3 correspond to variants from two different clades in the network. E. Simulated networks were compared to inferred networks based on output antibody sequences. Both correctly and incorrectly inferred networks are visualized. Each node depicts a unique variant, defined by a unique full-length VH-VL nucleotide sequence. The color of each node corresponds to a customizable repertoire feature (e.g., isotype, organ, phenotype) of cells within that variant of a certain isotype. Edges depict clonal relationships. Black nodes correspond to unmutated reference germline.

### Echidna simulates dynamic and time-resolved single-cell transcriptomes

During the time-resolved simulation of the immune receptors, each cell is additionally associated with a corresponding transcriptome containing gene expression information. The transcriptomes are simulated using either user-supplied distributions or transcriptional distributions based on publicly available single-cell immune repertoire sequencing data from 10x Genomics or recent publications (Bieberich *et al.*, 2021; Kuhn *et al.*, 2021; Neumeier, Pedrioli, *et al.*, 2021). It is additionally possible to supply a finite number of distinct transcriptional phenotypes and a corresponding vector, thereby dictating the average expression for each gene in each phenotype. At each time step, every cell will have a probability (either default or user-supplied) to transition into a new cell state. While under default conditions these are lineage-specific (e.g., a plasma cell will not revert back to a naive B cell), the user can override this transition matrix (in addition to each corresponding gene expression phenotypes) in the case modeling of custom cell populations is desired. To demonstrate the ability of Echidna to generate transcriptomes for adaptive immune cells, we simulated diverse CD8 T cell populations and their corresponding TCRs and transcriptomes, performed unsupervised clustering, and visualized them using uniform manifold approximation projection (UMAP) (Figure 4A). Overlaying user-supplied cell phenotype labels of naive, memory, effector, and exhausted T cells demonstrated that the cluster separation observed by UMAP corresponded to distinct cell phenotypes. One major strength of the simulation format of Echidna is that it is compatible with widely used scSeq analysis tools, such as Seurat (Satija *et al.*, 2015). Leveraging this compatibility, we demonstrated that the clusters indeed had distinct gene signatures corresponding to naive, memory, effector, and exhausted T cells via both violin plots and relating expression to location on the UMAP (Figures 4B, 4C).

**Figure 4.**
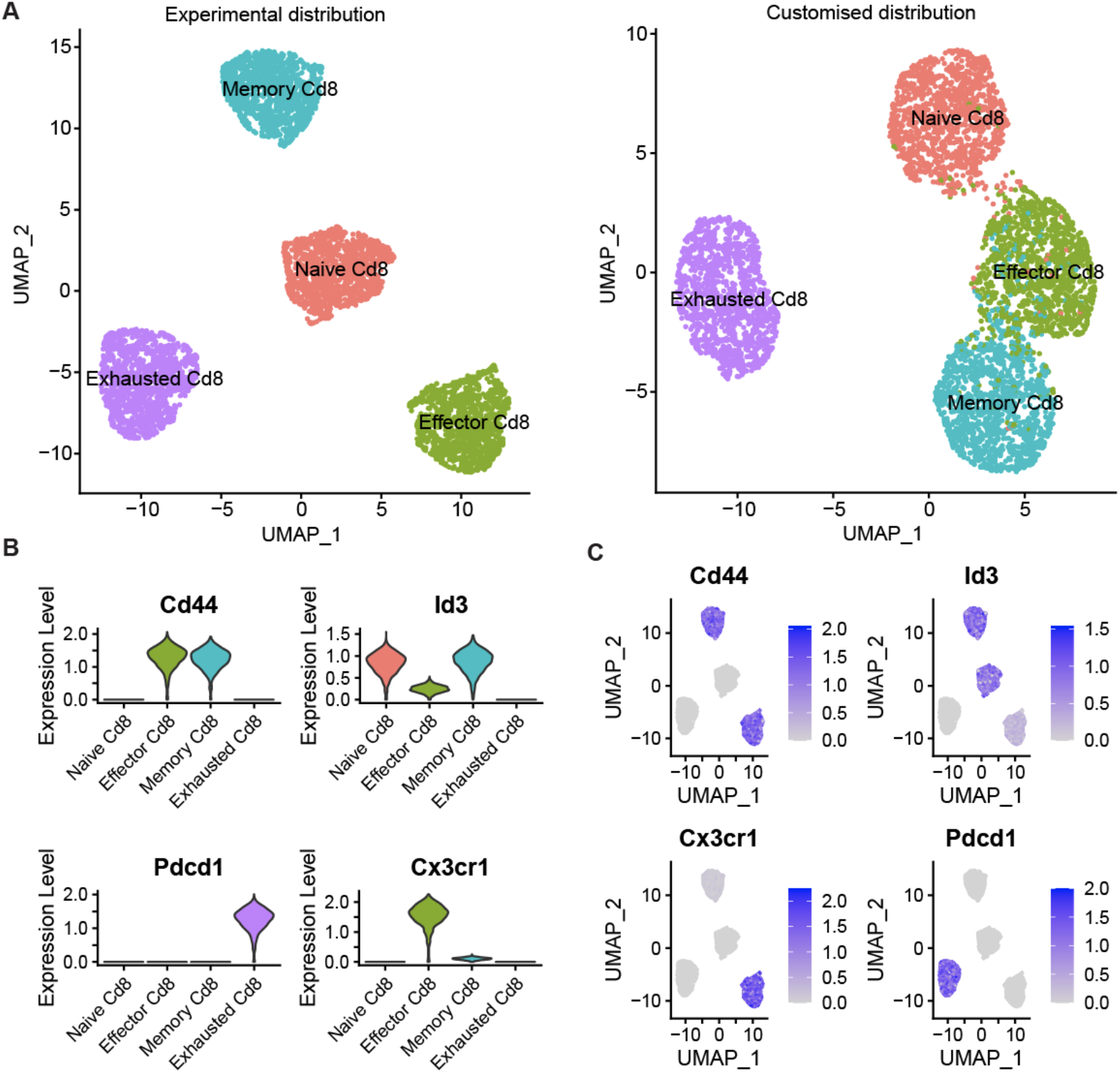
Echidna simulates transcriptomes based on experimental sequencing data. A. Uniform manifold approximation projection (UMAP) depicting transcriptional landscape of simulated CD8 T cell repertoires based on phenotypes from either experimental data or customized gene expression distributions. Color corresponds to clusters as determined by unsupervised clustering. Simulation output was used as input to the common scSeq tool Seurat. Each point represents a cell. B–C. Normalized expression of simulated T cells for select phenotype-defining genes visualized by either volcano plots or UMAP.

### Integrated simulation of immune receptors and transcriptomes

We lastly demonstrated the ability to integrate simulated adaptive immune receptors with gene expression information. We first generated BCRs and their corresponding transcriptomes at the single-cell resolution and quantified the cluster membership for the 50 most expanded B cell clones using parameter settings where certain cell phenotypes preferentially undergo clonal expansion (Figures 5A, 5B). The expanded clones could be visualized using UMAP, which revealed that clonally expanded cells were more likely to be in the cluster corresponding to the plasma cell phenotype relative to naive, germinal center (GC), and memory B cells (Figure 5C). This was further exemplified by quantifying normalized expression values of genes defining each of these subtypes, with plasma cell markers *Taci* and *Cd138 (Sdc1)* increased relative to markers characteristic of the other B cell subsets (*Cd19, Fas, Cd38)* (Figure 5D). In addition to seamlessly integrating simulated adaptive immune receptors into the popular R package Seurat, the transcriptional phenotypes and clusters can additionally be integrated into the mutational model, where certain antibody variants within a given clone can occupy distinct transcriptional states. To demonstrate this, we simulated two mutational networks, one where the transcriptional probability matrix was constant for all cells and another which had transcriptional probabilities specific for individual cells (Figure 5E), highlighting the ability of Echidna to simulate phenotype-driven SHM.

**Figure 5.**
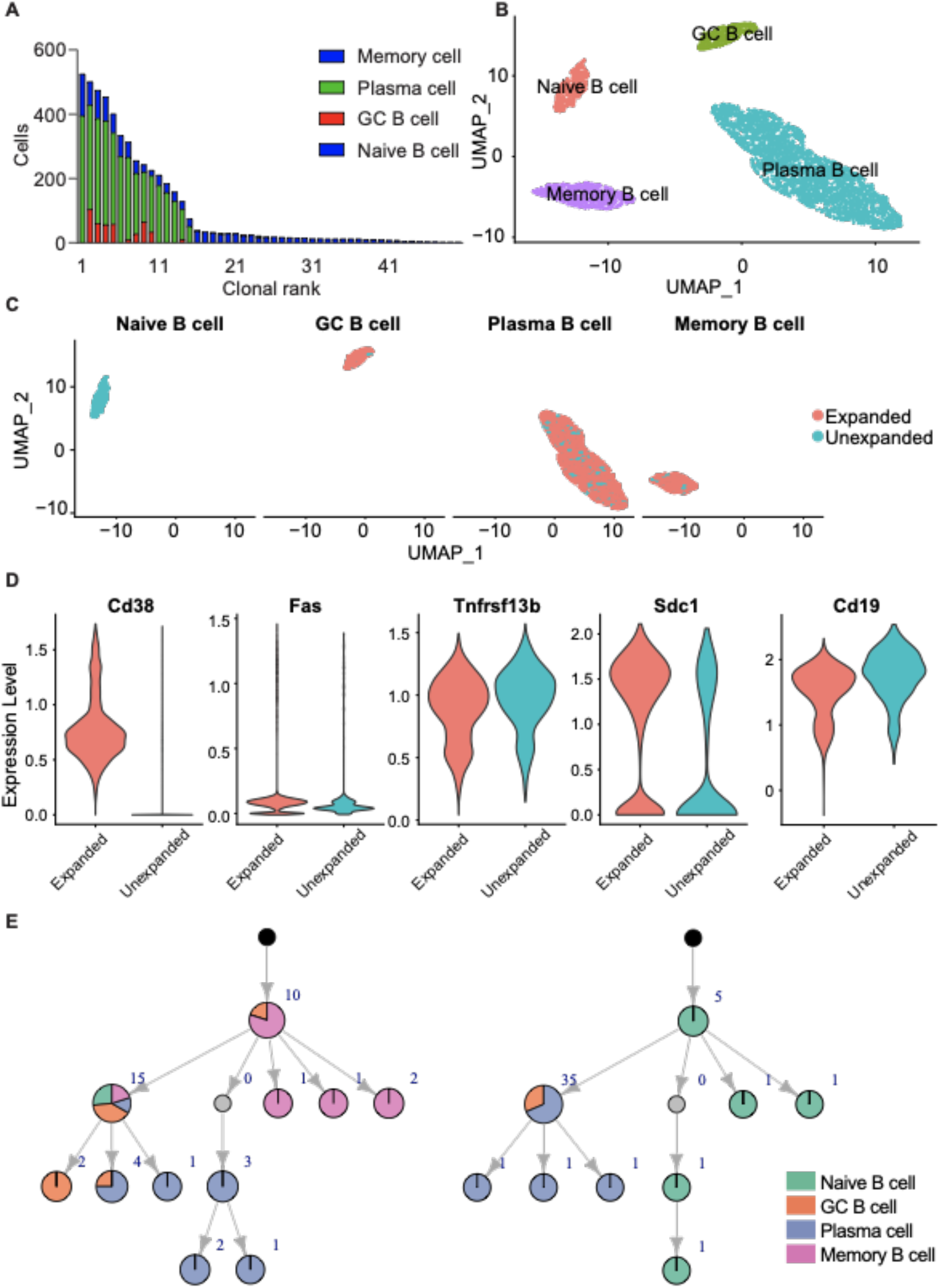
Echidna integrates gene expression and immune receptor during single-cell simulation. A. Preferential expansion of B cell subsets based on cell phenotype. Color corresponds to the number of cells within a given clone of a certain phenotype. Clone is defined as all cells belonging to a single simulated VDJ recombination event. B. Uniform manifold approximation projection (UMAP) highlighting distinct transcriptional profiles of naive, germinal center (GC), memory, and plasma cells. C. UMAP highlighting expanded and unexpanded clones. Expanded clones are defined as those clones supported by two or more unique cells. D. Normalized expression of *Cd38*, *Fas*, *Taci*, *Cd138*, and *Cd19* separated by expanded vs unexpanded cells. E. Simulated clonal lineage depicting that certain B cell subtypes can be customized to preferentially undergo somatic hypermutation or clonal expansion.

## Discussion

Recent technological advancements in single-cell immune repertoire sequencing have provided insight into how clonal selection relates to cellular phenotypes, thereby paving the way to simultaneously model gene expression with adaptive immune receptor sequences. Here, we developed a simulation framework and further exemplified how simulated repertoires can be leveraged to benchmark and develop bioinformatics software relating to experimental datasets. Although multiple simulation frameworks have been previously developed (Yermanos *et al.*, 2017; Weber *et al.*, 2020; Safonova *et al.*, 2015; Davidsen and Matsen, 2018), there is a lack of software specifically tailored to generating adaptive immune receptors at the single-cell resolution and with corresponding transcriptome information. Echidna bridges the gap between single-cell transcriptome and immune repertoire sequencing, providing information relevant to both fields at the single-cell resolution. An additional advantage of Echidna is that mutations are introduced at the clonal level in a time-resolved manner while also providing the true, known network structure that generated a given clonal lineage. This information can be used to explore how various parameters dictating inference of phylogenetic trees and similarity networks relate to either the immune repertoire or gene expression phenotypes (DeWitt *et al.*, 2018). Although we have provided default parameter values derived from experimental sequencing data from 10x Genomics and recently published data (Kuhn *et al.*, 2021; Bieberich *et al.*, 2021; Neumeier, Pedrioli, *et al.*, 2021). Echidna similarly allows the user to customize each feature of the simulation, including the initial clonal diversity generated via V(D)J recombination, clonal expansion, transcriptional states, and their respective transition probabilities, and, for B cells, both SHM and isotype distributions.

Modeling immunological processes requires significant approximations and assumptions that may not accurately reflect reality. Congruent with this, our simulation tool relies upon many assumptions at each step in the algorithm. As an example, the default simulation utilizes a simple method to simulate V(D)J recombination that relies upon selecting germline genes and inserting/deleting nucleotides based on subjective probability distributions. While this produces extremely diverse repertoires, it ignores many of the complexities not immediately observable underlying actual repertoire diversity (Marcou *et al.*, 2018; Greiff *et al.*, 2017; Slabodkin *et al.*, 2021). We, therefore, have introduced the ability for more sophisticated repertoire simulations involving VAEs (Davidsen *et al.*, 2019; Friedensohn *et al.*, 2020), which under default parameters were trained on publicly available data of naive B and T cell repertoires. Importantly, we have additionally included the option for users to supply their own repertoires as training data to the VAE, thereby enabling the simulation of a repertoire resembling specialized experimental conditions. As more datasets linking gene expression and immune receptors become publicly available, we can continuously update the generative model underlying our framework, thereby bringing our simulations closer to the generative model underlying natural immune repertoires.

## Methods

### Simulating adaptive immune repertoires at single-cell resolution

Relevant simulation parameters for all figures can be found as supplementary information (Table S1) and accompanying code and compiled package can be found at github.com/alexyermanos/echidna. The VDJ recombination simulation portion of the algorithm is adapted from AbSim (Yermanos *et al.*, 2017). Here, we simulate the heavy chain and light chain sequence of an individual cell simultaneously and integrate the relevant sequence and annotation information. For heavy chain simulations, D and J gene segments are first combined, followed by V segments joining to the previously formed D-J segment. In the case of light chains, only V and J segments are joined. Insertions and deletions can occur at the junction site for each fusion event. In the case of insertions, each of the four nucleotides had an equal probability of being selected. The number of inserted nucleotides can range from 0 to 10 under default parameters but can either follow a uniform or a custom probability distribution supplied from an experimental human or mouse-specific model, as specified by the user. For junctional deletions, the sampled number of deleted nucleotides at the end of each germline segment can be customized. However, for all simulations presented here, a range of 0 to 5 nucleotides was selected with a uniform distribution. The V, D, and J segments were sampled using a uniform distribution from all human and mouse germline genes from IMGT (Lefranc *et al.*, 2003), unless otherwise specified. The user can modify their frequency by adding, replicating, or deleting genes in the reference germline genes. For the example presented in this manuscript, IGHV1-11 and IGKV1-131 genes were replicated in the input germline gene list to obtain preferential heavy and light chain gene usage.

The random insertion and deletion processes in the VDJ recombination mechanism might introduce frameshift mutations and generate non-productive sequences before any evolution commences. We have therefore provided an option for limiting the initial repertoire to only productive sequences as determined by using the align and exportAlignments commands from MiXCR (v3.0.1) under default parameters (Bolotin *et al.*, 2015). After restricting the initial VDJ recombination events to productive sequences, the repertoire will only use these sequences to initialize new clones. The productive sequences were prepared by first simulating more than 100,000 VDJ sequences for each chain type and species and subsequently filtering out non-productive sequences with MiXCR (Bolotin *et al.*, 2015) from simulated sequences. The productive sequences were additionally annotated with their CDR3 nucleotide and amino acid sequences.

### Simulating variational autoencoders

As an additional option to simulate productive adaptive immune receptors, we employed variational autoencoders (VAEs) to generate new sequences with patterns resembling experimental data (Friedensohn *et al.*, 2020; Davidsen *et al.*, 2019; Eguchi *et al.*, 2020) using R package Keras (JJ Allaire and François Chollet 2020). Our VAE pipeline was based on the example script in the variational_autoencoder function of the Keras package. VAE models were trained based on publicly available repertoire sequencing data (Table S2). Input data sets were first one-hot encoded and subsequently split into two evenly sized folds. One fold was used for validation and the remaining folds were used for training, where the maximum range of latent points was then recorded after model training. For each chain type, 1,000,000 points were drawn within the recorded range in latent space containing the training data (Figure S3). The predictive values of decoded sample points were then reconstructed into nucleotide sequences based on the highest predicted value of the VAE model. A threshold of the four predicted values for the four nucleotide bases was set to determine the end of the sequence. The base was considered to be null if the predicted value was lower than the threshold (0.05). Final sequences were generated from the generator model using the predict function from stats R package with the generative object supplied as input under default parameters. The final VDJ sequences were then filtered by MiXCR to ensure only productive sequences as previously described, which were embedded in the starting repertoire of sequences for subsequent simulation.

### Somatic Hypermutation

The method to simulate SHM for B cells is adapted from AbSim (Yermanos *et al.*, 2017), which offers multiple methods to introduce sequence diversity. First, a “Poisson” method can mutate each nucleotide randomly at a user-defined probability at every position in the immune receptor. Second, the “Data-driven” method involves targeted mutations in inferred CDR locations, in addition to allowing different mutation rates for nucleotide transitions or transversions. Third, the “motif” method considers the influence of neighboring bases and detects 5-mer sequence motifs in the sequence and applies nucleotide-specific substitution probabilities to nucleotides located in the middle of 5-mer motifs. The transition probabilities were determined from previous studies modeling mutational frequencies across multiple antibody datasets (Yaari *et al.*, 2013). If desired, the user can additionally update the 5-mer mutational frequencies with custom distributions. Additionally, a “wrc” method is available, where the WRC motifs in the antibody sequence will preferentially undergo somatic hypermutation substitution based on the relevant 5-mer motifs (Yaari *et al.*, 2013). A combination of any of the above-mentioned methods with user-defined weights is possible. Finally, the user can specify custom SHM probabilities for certain repertoire features, such as transcriptional phenotypes and antibody isotypes.

### Simulated gene expression

The gene expression level of a certain defined phenotype can be represented with a numeric base vector, where each element corresponds to the normalized expression level for each gene. The final expression level for an individual cell will be sampled based on the base vector in combination with a user-defined noise parameter. The noise distribution can be different for each specified transcriptional phenotype. Therefore, cells of the same phenotype will sample from an identical base vector, but their actual value of expression level will fluctuate with the introduction of noise. Finally, gene expression vectors will be combined into a gene expression matrix that is compatible with common RNA-seq analysis frameworks, such as Seurat (Satija *et al.*, 2015). For B cells, the user can choose any combination of the four phenotype switching strategies by enabling or disabling any of them: (i) SHM dependent: If SHM occurs to a cell, it will switch to one of the user-supplied possible phenotypes. If there is more than one potential phenotype, the probability to change from one phenotype to the next is determined by the transition matrix. In the default transition matrix, the chance of GC B cells switching to plasma cells and memory B cells are 2/3 and 1/3, respectively. Whereas, under default parameters, plasma cells would not transition to other phenotypes when undergoing SHM, as their transition probabilities to the other cell types are set to 0. (ii) Class switching dependency: Similarly, phenotype switching can occur once a cell undergoes class-switch recombination, again defined by a transition matrix containing the probabilities to change from one state to another state. (iii) Variant selection dependency: within each clone, an individual variant (or node in the network) will have a chance to undergo phenotypic transition. Therefore, certain B cell sequences (e.g., certain affinities) can be modeled to adapt novel phenotypes based on antibody sequence. (iv) Phenotypes switch randomly following either a default transition matrix or user-supplied transition matrix.

To generate default gene expression vectors, we incorporated data from either 10x Genomics or previous single-cell immune repertoire sequencing studies containing either human or mouse B and T cells (Table S3) (Bieberich *et al.*, 2021; Kuhn *et al.*, 2021; Neumeier, Pedrioli, *et al.*, 2021). The data was prepared by first pooling all cells (separately for human and mice) and subsequently sorting cells of desired phenotypic markers. The average gene expression levels across all cells matching the phenotypic marker definitions were then extracted as base vectors for each phenotype (Table S3). While these serve as default parameters, the package is intended to receive custom expression vectors based on a user’s individual dataset or interest.

### Inferring mutational networks

Mutational networks were inferred as previously described (Neumeier, Pedrioli, *et al.*, 2021), which utilizes the R package igraph (Csardi *et al.*, 2006). First, the pairwise edit distance was calculated for each appended, full-length heavy and light chain sequence (including the reference germline), thereby generating a distance matrix for each clone using the stringdist R package (van der Loo, 2014). This distance matrix was then used to determine the order in which sequences were added to the network. The unmutated germline reference gene initializes each network, and then the sequence with the smallest distance was added to the most similar sequence in the network in an iterative manner. Edit distance ties were resolved by randomly selecting from the potential nodes within the network. The reference germline sequence and appended heavy and light sequence of each clone are available in the simulation output.

## Data Visualization

Circos plots were created using the function VDJ_circos in Platypus (v3.1), which relies upon the R package circlize (v0.4.13) (Gu *et al.*, 2014). Mutational networks were created using igraph (v1.2) (Csardi *et al.*, 2006). Sequence alignment was performed in Geneious Prime (v.2020.0.3) under default parameters. UMAP, violin plots, and feature plots were performed using Seurat (v4.0.1) (Satija *et al.*, 2015). Remaining plots were produced in Prism Graphpad (v9).

## Data availability

The R package and code used in this manuscript can be found at github.com/alexyermanos/echidna and also in the R package Platypus (Yermanos, Agrafiotis, *et al.*, 2021b). Publicly available data and corresponding sample accession numbers can be found Supplementary Tables S2 and S3.

## Supporting information for

**Figure S1.**
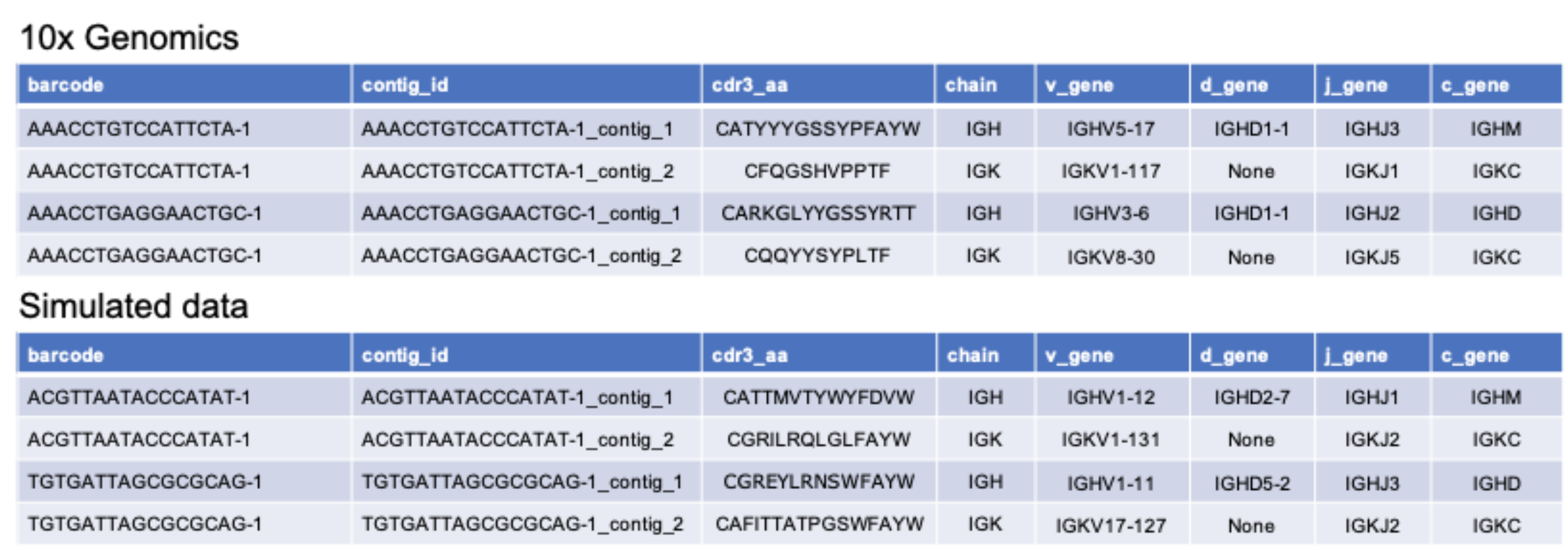
Simulated repertoire output mirrors 10x Genomics’ cellranger output from experimental sequencing datasets.

**Figure S2.**
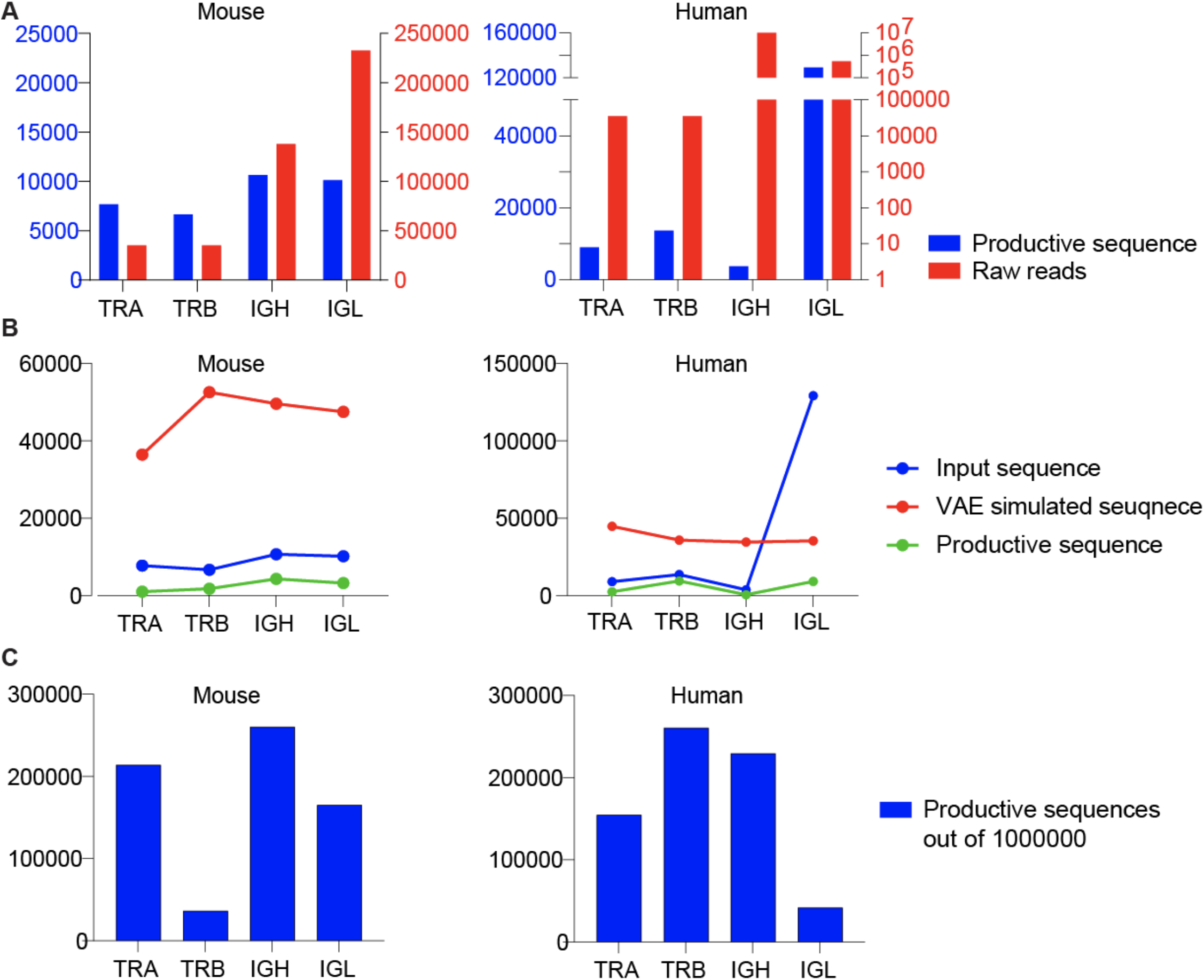
Fraction of productive V(D)J recombination events. A. The number of raw reads and the number of productive adaptive immune receptor sequences per variable chain for human and mice from publicly available data. B. The number of productive input sequences used as input to train the variational autoencoder (VAE), the number of simulated sequences by the VAE generative model, and the number of productive simulated sequences as determined by MiXCR. C. The number of productive sequences out of 1,000,000 simulated VDJ recombination events using a naive model of appended reference alleles from IMGT together with uniform probabilities of insertions and deletions at each simulated junction.

**Figure S3.**
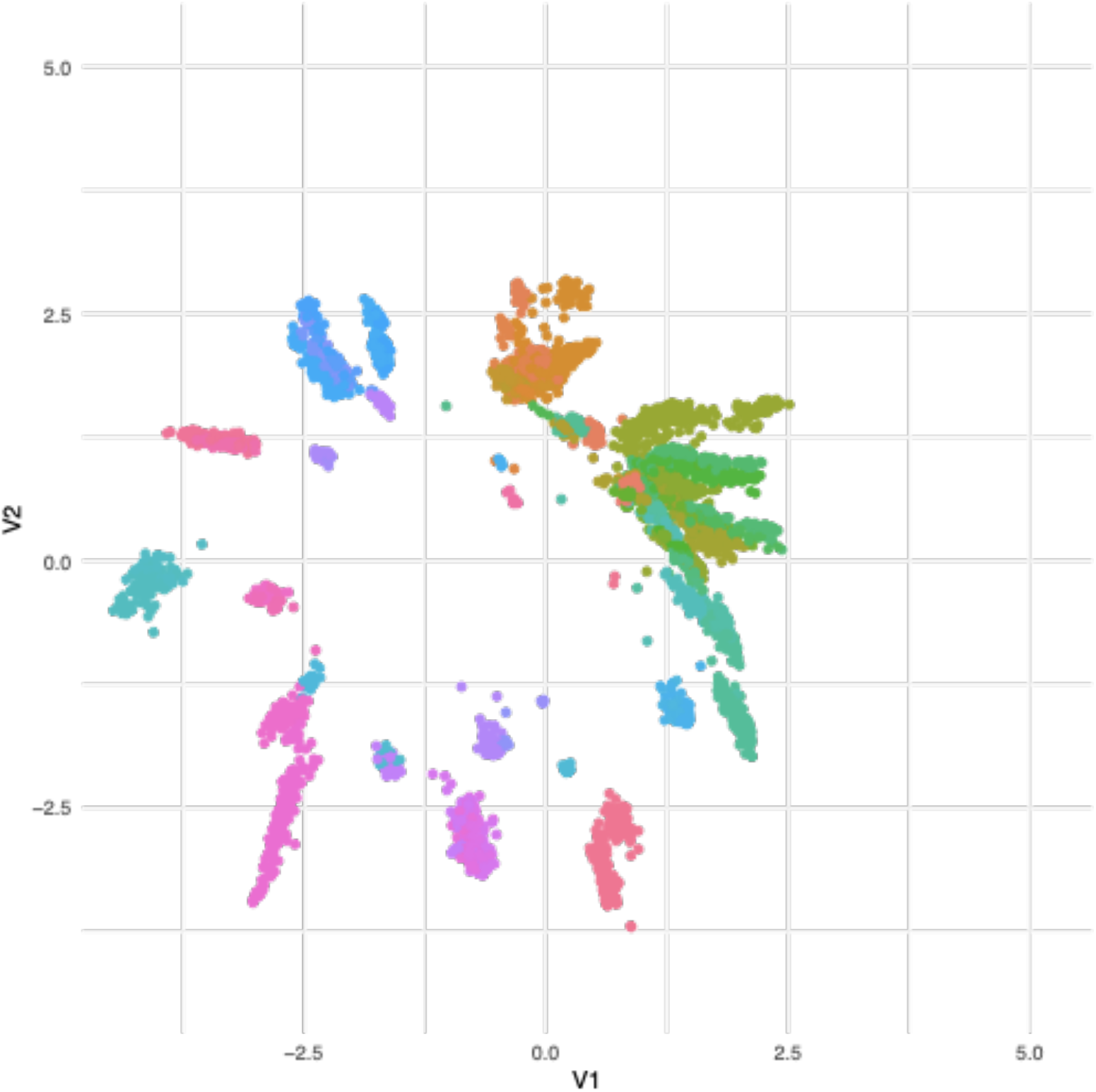
Example latent space of the murine heavy chain variational autoencoder (VAE). Color corresponds to a unique germline V gene segment. Full-length variable region (framework region 1 to framework region 4) was used as input to the VAE.

**Figure S4.**
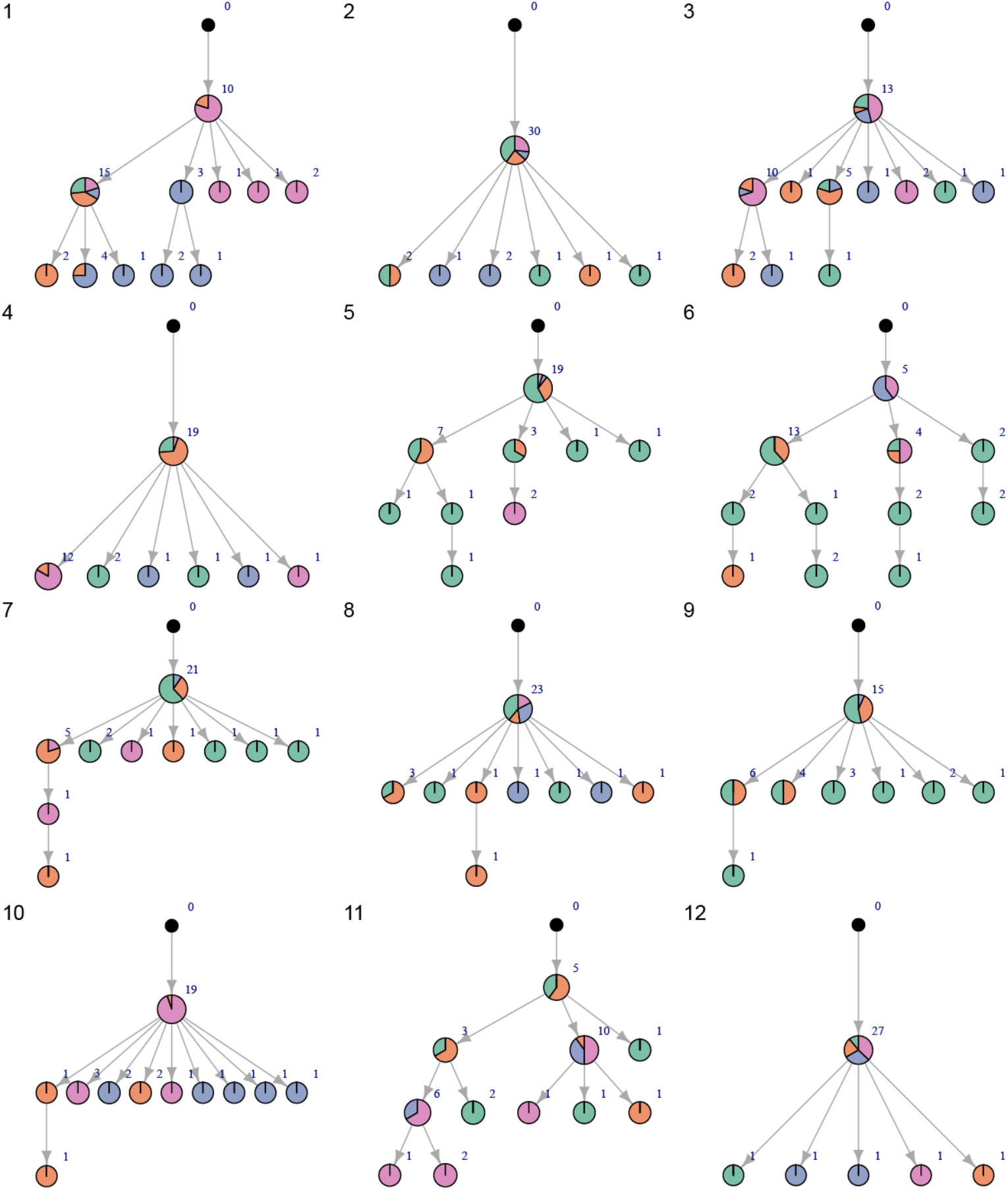
Examples of correctly inferred networks based on output sequences from Echidna. Color corresponds to user-tunable cell phenotype parameters. Nodes correspond to unique, full-length nucleotide variants. Node label corresponds to the number of cells with an identical antibody sequence.

